# Replication Timing and Transcription Identifies a Novel Fragility Signature Under Replication Stress

**DOI:** 10.1101/716951

**Authors:** Dan Sarni, Takayo Sasaki, Karin Miron, Michal Irony Tur-Sinai, Juan Carlos Rivera-Mulia, Brian Magnuson, Mats Lungman, David M. Gilbert, Batsheva Kerem

## Abstract

Common fragile sties (CFSs) are regions susceptible to replication stress and are hotspots for chromosomal instability in cancer. Several features characterizing CFSs have been associated with their instability, however, these features are prevalent across the genome and do not account for all known CFSs. Therefore, the molecular mechanism underlying CFS instability remains unclear. Here, we explored the transcriptional profile and temporal order of DNA replication (replication timing, RT) of cells under replication stress conditions. We show that the RT of only a small portion of the genome is affected by replication stress, and that CFSs are enriched for delayed RT. We identified a signature for chromosomal fragility, comprised of replication stress-induced delay in RT of early/mid S-phase replicating regions within actively transcribed large genes. This fragility signature enabled precise mapping of the core fragility region. Furthermore, the signature enabled the identification of novel fragile sites that were not detected cytogenetically, highlighting the improved sensitivity of our approach for identifying fragile sites. Altogether, this study reveals a link between altered DNA replication and transcription of large genes underlying the mechanism of CFS expression. Thus, investigating the RT and transcriptional changes in cancer may contribute to the understanding of mechanisms promoting genomic instability in cancer.

## Introduction

DNA replication occurs in a temporal order known as the replication timing (RT) program, which is tightly regulated to ensure the faithful duplication of the genome (Rivera-Mulia and Gilbert, 2016). DNA replication stress leads to formation of DNA damage and subsequently to genomic instability (Alexander and Orr-Weaver, 2016; Gaillard et al., 2015; Macheret and Halazonetis, 2015; Zeman and Cimprich, 2014). Replication stress induced by DNA polymerase inhibitors or activation of oncogenes leads to decreased replication fork progression, stalled forks, and activation of dormant origins to complete duplication of the genome (Alver et al., 2014; Ge et al., 2007; Halazonetis et al., 2008; Macheret and Halazonetis, 2015). However, activation of dormant origins is not always sufficient and thus such perturbed replication induces chromosomal instability at specific regions termed common fragile sites (CFSs) (Glover et al., 1984). CFSs manifest as breaks, gaps and constrictions on metaphase chromosomes under mild replication stress (Glover et al., 2017). CFSs are preferentially unstable in pre-cancerous lesions and during cancer development implying that they play a role in driving cancer development (Bartkova et al., 2005; Bignell et al., 2010; Gorgoulis et al., 2005; Tsantoulis et al., 2008). In fact, several fragile sites were found within tumor suppressor genes such as *FHIT* and *WWOX* that are unstable in a wide variety of different cancer types (Gao and Smith, 2015; Karras et al., 2016).

Several features characterizing CFSs have been proposed to contribute to CFS expression (fragility), including late replication (Le Beau, 1998; Letessier et al., 2011) and a paucity of replication origins (Letessier et al., 2011), which under replication stress impede the completion of their replication. DNA secondary structures induced by AT-rich sequences have also been suggested to cause CFS expression by raising potential barriers to replicating forks (Shah et al., 2010; Zlotorynski et al., 2003). Additionally, CFSs are enriched with large genes (McAvoy et al., 2007) that may contribute to fragility due to replication-transcription collisions (Gaillard and Aguilera, 2016; Helmrich et al., 2011; Sollier and Cimprich, 2015). However, none of the suggested factors on their own is sufficient to induce fragility since they are prevalent across the genome including non-fragile regions. Furthermore, the suggested fragile site features could not account for all known CFSs (Sarni and Kerem, 2016). Thus, the molecular basis underlying recurrent chromosomal instability remains unknown.

CFS instability was found to be both cell type- and stress inducer-specific, raising the possibility that fragility is driven by perturbed regulation of organized yet dynamic cellular programs, such as DNA replication and transcription. Therefore, we hypothesized that a combination of several factors together render a region sensitive to replication stress and induce fragility. In order to investigate this premise we tested the effect of mild replication stress induced by the DNA polymerase inhibitor aphidicolin (APH) on genome-wide replication and transcription programs, using Repli-seq and Bru-seq, respectively.

Here, we have identified a signature for chromosomal fragility upon replication stress, comprised of APH-induced delay in RT of early/mid-S phase replicating regions and active transcription. We show that the temporal order of replication of only a small part of the genome is altered by APH, and the delayed portion is highly enriched for CFSs. The induced replication delay generates a V-shaped profile, which in CFSs is associated with transcribed large genes. Moreover, we show that the CFS core fragility regions in unperturbed cells are replicating in mid S-phase and thus are not merely the latest replicating regions as previously suggested (Le Beau, 1998; Letessier et al., 2011). Furthermore, our “fragility signature” enabled greater precision of mapping the core fragility region, the identification of novel fragile sites, not detected cytogenetically, highlighting the improved sensitivity of our approach for identifying fragile sites. Altogether, the combined analysis of RT and transcriptional profiles under replication stress reveals a link between altered DNA replication and transcription of large genes underlying CFS expression.

## Results

### Aphidicolin affects the replication timing of specific genomic regions

To study the effect of replication perturbation on the replication program in general and at CFSs in particular, we performed Repli-seq on telomerase-immortalized human foreskin fibroblasts (BJ-hTERT). Unchallenged cells were compared to cells treated with a low concentration (0.2 μM) of APH for 24h (control and APH, respectively), commonly used to induce CFS expression (Figure 1a), thus cells were exposed to the APH for at least one cell cycle prior to analysis. Genome-wide correlation analysis of RT profiles was able to distinguish control from APH-treated cells (Figure 1b), despite a strong genome-wide correlation of RT profiles among all samples (Figure 1b). Next, we extracted all RT-variable regions by segmenting the genome into 100kb windows and applied an unsupervised K-means clustering analysis to all 100kb genomic regions that changed their RT by at least 1 RT unit (Log2 ratio of early/late), as previously described (Rivera-Mullia et al., 2015). We identified 1,277 RT-variable regions (4% of the genome), consistent with the correlation matrix analysis (Figure 1b,c). Many RT-variable 100kb windows were contiguous and, when assembled together, identified 607 regions differentially replicated upon APH treatment. Clustering the RT-variable regions into 5 clusters identified specific RT signatures containing early replicating regions in control cells that are delayed in APH treated cells (RT signature 3) and regions that replicate late in control cells but their replication was advanced in APH treated cells (RT signature 1) (Figure 1c,d). Moreover, we identified regions where RT changed within the same fraction of S-phase (early or late) following APH treatment, e.g. regions that replicate very early in control cells and are delayed by APH treatment yet still replicate in early S-phase in APH treated cells (RT signature 4) and vice versa (RT signature 5) (Figure 1c,d). Examples of RT signature profiles showing alterations following APH treatment are shown in Figure 1d. It is worth noting that 2/3 of RT-variable regions (387) were delayed by APH treatment (Supplementary Table 1). These delayed loci are of interest for further investigation into chromosomal stability under replication stress and in particular at CFSs, as mild replication stress was shown to induce under-replication associated with CFS expression (Le Beau, 1998; Minocherhomji et al., 2015).

**Figure 1.**
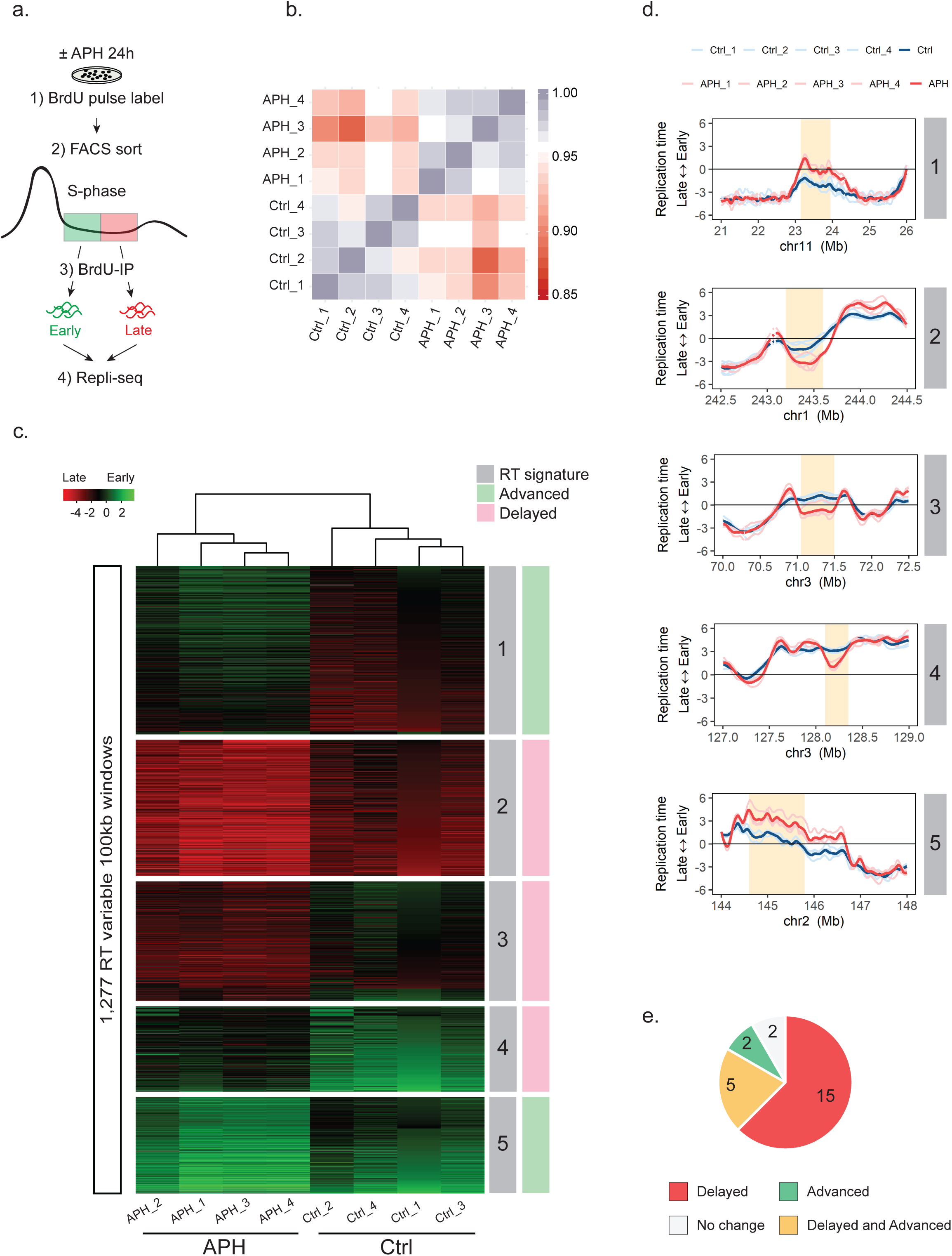
Aphidicolin affects the replication timing of specific genomic regions. (**a**) Schematic description of Repli-seq. BJ-hTERT cells with (+) or without (-) APH treatment were pulse labeled with BrdU and sorted into early and late S-phase, and the RT programs were obtained by next-generation sequencing. (**b**) Correlation matrix of genome-wide RT programs of BJ-hTERT cells with (APH) or without (Ctrl) APH treatment. 4 replicates per condition are presented (1-4). (**c**) Identification of APH-specific RT signatures, distinguishing APH-treated cells (APH) from non-treated control cells (Ctrl). Unsupervised k-means clustering analysis of RT-variable regions identified specific RT signatures (labeled in gray boxes). The heat map shows the RT ratios [= log2(early/late)]. (**d**) Exemplary RT profiles of RT signatures. Numbering according to RT signature in (**c**). The RT profiles are displayed as log2 ratios of signals from early and late S-phase fractions. Positive values correspond to early replication, and negative values correspond to late replication. Four replicates per condition are presented color coded according to the legend (Ctrl in blue, APH in red). Yellow boxes mark the RT variable regions. (**e**) Pie chart indicating RT alterations within CFSs mapped in BJ-hTERT cells following APH treatment.

To test whether CFSs are enriched with RT-variable regions we analyzed the RT of the CFSs we have previously identified in BJ-hTERT cells (Miron et al., 2015). The analysis showed delayed replicating loci in 20/24 (83%) of identified CFSs in cells treated with APH relative to the control cells (Figure 1e). Advanced-RT regions were found only in 7/24 (29%) CFSs, amongst them 5 CFSs harbored both advanced and delayed RT loci (Figure 1e). These results indicate that CFSs are enriched for delayed-RT. Next, we trisected the RT program into early, mid and late S-phase according to the RT value. We then compared the timing of replication for the APH-induced RT delayed regions in control cells revealing that 80% of the delayed-RT loci in CFSs are replicated in early/mid-S phase. Thus, the APH-induced RT-delayed loci are not the latest replicating regions in unchallenged cells.

### Delayed replication of large genes at CFSs is associated with fragility

Cytogenetic mapping enables the identification of fragile sites on metaphase chromosomes of cells grown under replication stress conditions. However, due to its low resolution (Mb in size), cytogenetic mapping does not allow the identification of the core fragility region within a cytogenetic band harboring a CFS. In order to investigate whether the delayed replication in APH treated cells is the fragility core within CFSs we used molecular mapping using BAC clones as fluorescence *in-situ* hybridization (FISH) probes on metaphase chromosomes from cells treated with APH. We first focused on two CFSs that we have previously identified in BJ-hTERT cells (Miron et al., 2015); the first located on chromosome 2q22 (2q22) and the second on chromosome 22q12 (22q12).

The cytogenetic 2q22 band spans over 10Mb and has three prominent late replicating regions in control cells (Figure 2a), which do not change following the APH treatment (marked in grey, Figure 2a) and one mid-S replicating plateau region that is substantially delayed following APH treatment (marker in orange, Figure 2a). Interestingly, this RT-delayed domain coincides with a large gene, *GTDC1* (386 kb, Figure 2a). Moreover, flanking *GTDC1* are two noticeable large mid-S replicating domains in control cells, which replicate earlier following APH treatment (Figure 2a). Thus, APH treatment induced a V-shaped RT pattern at the *GTDC1* locus, which indicates a lack of origin firing within the gene body under replication stress and earlier activation of origins flanking the delayed region. Additionally, the combination of a delayed replication region flanked by advanced replication generated steep RT slopes, indicating that under replication stress the locus is replicated by single/few slow and long traveling replication forks. This is in contrast to the RT plateau at the *GTDC1* locus in control cells, implying that under normal conditions the speed of replication fork progression is sufficient to properly replicate the gene. Therefore, the APH-induced V-shaped RT profile may represent a high resolution signature of fragility.

**Figure 2.**
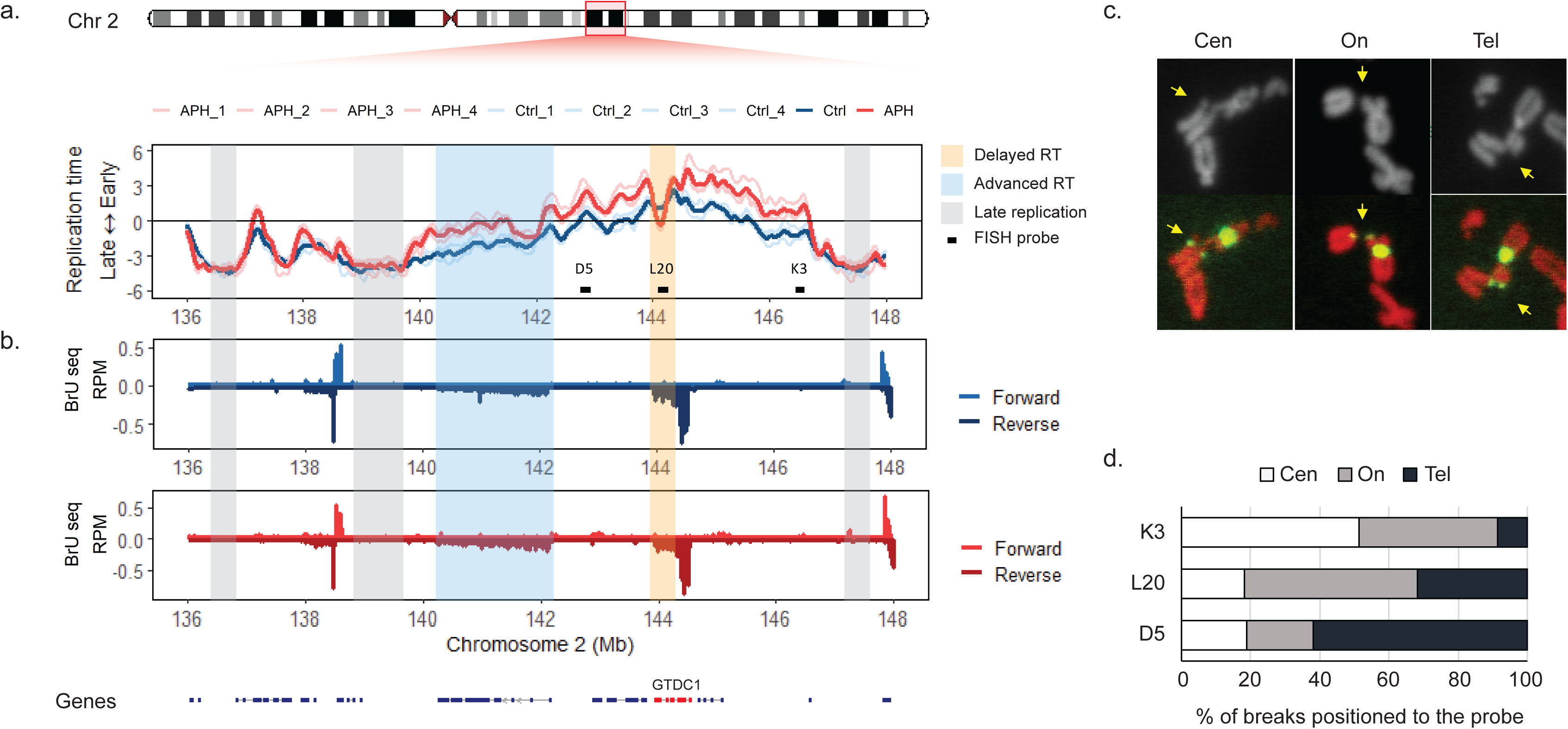
Delayed replication of *GTDC1* at the CFS 2q22 is associated with fragility. (**a**) RT profile of the fragile site 2q22 showing advanced RT flanking delayed replication of *GTDC1*. Grey boxes mark the latest replicating regions within the 2q22; Blue box marks the non-fragile *LRP1* gene; Yellow box marks the fragility core within *GTDC1*. FISH probes are presented as black boxes. Four replicates per condition are presented color coded according to the legend (Ctrl in blue, APH in red). (**b**) Nascent RNA transcription (nRNA-seq), Bru-seq, traces for Ctrl and APH treated cells at 2q22. Positive values represent forward strand, negative values represent reverse strand. One replicate per condition is presented, color coded: Ctrl in blue, APH in red. Annotated genes are shown at the bottom. *GTDC1* is label in red. (**c**) Representative images of chromosomal breaks on metaphase spreads labeled with a FISH probe, L20, marked in green. DNA stained with PI marked in red. Yellow arrows indicate break points. Three types of breaks presented positioned relative to the probe: Closer to the centromere (Cen), closer to the telomere (Tel) and a split probe signal (On). (**d**) Quantification of chromosomal aberrations positioned relative to the probe, as represented in (**c**), showing a shift in breaks from K3 to D5, indicating the fragility core in between these probes.

To test whether the APH-induced V-shaped RT profile is indeed the site of fragility we next mapped the core fragility region within 2q22 using FISH mapping. We used fluorescently labeled probes (D5, L20 and K3) delimiting the APH-induced V-shaped RT domain. Analysis of metaphase spreads hybridized with fluorescently labeled probes showed that breaks occurred at the replication stress sensitive V-shaped RT domain (Figure 2c,d), indicating that this locus within 2q22 is indeed the core fragility region of the fragile site. Moreover, the FISH analysis showed that the three prominent late replicating regions in control cells, which do not change following the APH treatment, are excluded from the fragility core (Figure 2). These results demonstrate that, the core fragility region is not located within the latest replicating regions at a CFS, but it lies within regions that under normal conditions replicate in early/mid S-phase and are delayed upon APH treatment.

To validate our hypothesis that APH-induced delayed RT is prone to fragility under replication stress, we mapped the core fragility region within a second CFS located at chromosome 22q12. This fragile band spans over 10Mb and has a prominent late replicating domain in control cells, which does not change following APH treatment (marked in grey, Figure 3a). Comparing the RT profiles of control and APH treated cells identified one noticeable domain of delayed replication following APH treatment, which in control cells is replicating in early/mid-S (marked in orange, Figure 3a). This domain coincides with a large gene, *TTC28* (∼700kb). Thus, the APH-induced delayed replication of the *TTC28* locus generated a novel replication stress associated V-shaped RT domain. It is worth noting that the delayed replication domain was not flanked by earlier replicating DNA in this case. Thus, the results indicate that the replication of the locus is susceptible to replication perturbation resulting in APH-induced delayed RT, prone to fragility under replication stress. Next, we analyzed metaphase spreads of APH-treated cells hybridized with FISH probes delimiting *TTC28*. The analysis showed that that the fragile site breaks and gaps occur between the FISH probes indicating that *TTC28* is the fragility core region within this CFS (Figure 3c,d). Thus, RT analysis and molecular mapping of the fragile sites 2q22 and 22q12 suggest that APH-induced replication delay is underlying CFS expression. Moreover, this delayed replication locus coincides with large genes suggesting that under replication stress they are replicated by single slow moving, aphidicolin-delayed, and long traveling forks, likely because the gene body is lacking dormant replication origins (pre-RCs) that it can recruit to accelerate replication. Interestingly, in both fragile sites we have molecularly mapped, the very late replicating regions, which do not change in RT, are outside of the core fragility region (Figure 2,3), suggesting that fragility is induced at regions in which RT is altered by replication stress and not at regions with stable RT, even if they are the latest replicating domains as previously suggested (Letessier et al., 2011). These results re-emphasize the importance of studying RT at CFSs under replication stress conditions.

**Figure 3.**
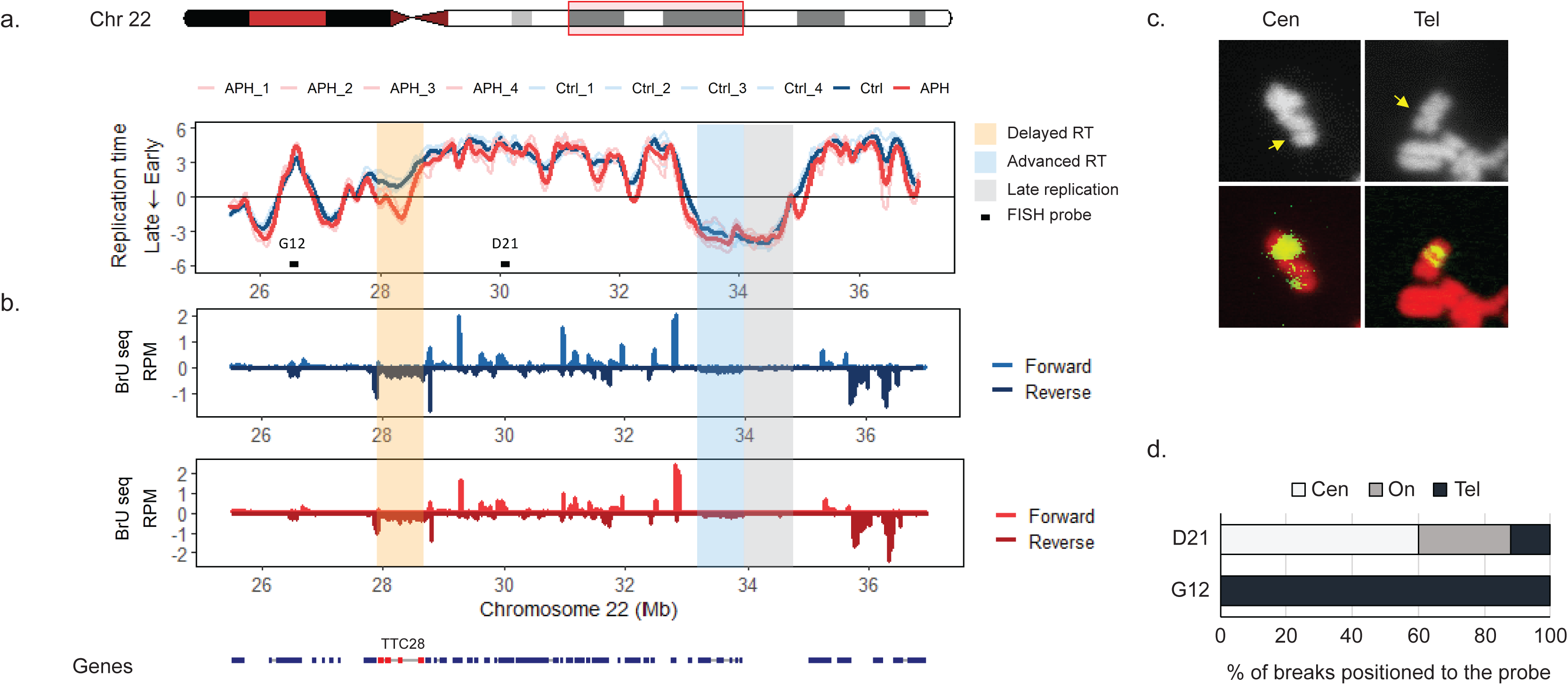
Delayed replication of *TTC28* at the CFS 22q12 is associated with fragility. (**a**) RT profile of the fragile site 22q12 showing advanced RT flanking delayed replication of *TTC28*. Grey box marks the latest replicating regions within the 22q12; Blue box marks the non-fragile *LARGE1* gene; Yellow box marks the fragility core within *TTC28*. FISH probes are presented as black boxes. Four replicates per condition are presented color coded according to the legend (Ctrl in blue, APH in red). (**b**) nRNA-seq (Bru-seq) traces for Ctrl and APH treated cells at 22q12. Positive values represent forward strand, negative values represent reverse strand. One replicate per condition is presented, color coded: Ctrl in blue, APH in red. Annotated genes are shown at the bottom. *TTC28* is label in red. (**c**) Representative images of chromosomal breaks on metaphase spreads labeled with FISH probes, G12 and D21, marked in green. DNA stained with PI marked in red. Yellow arrows indicate break points. Two types of breaks presented positioned relative to the probe: Closer to the centromere (Cen) and closer to the telomere (Tel). (**d**) Quantification of chromosomal aberrations positioned relative to the probe, as represented in (**c**), showing a shift in breaks from G12 to D21, indicating the fragility core in between these probes.

### Delayed replication of large genes is insufficient to drive CFS expression

CFSs were found to be enriched with large genes (>300 kb) (Le Tallec et al., 2013). Therefore, to investigate the association we observed between large genes, APH-induced RT delay and chromosomal fragility (Figure 2,3), we analyzed the RT profiles of all the genes in the genome (excluding genes <5 kb). RT analysis showed that genes >230 kb tend to replicate later in S-phase as compared with the rest of the genes in control cells (Figure 4a, Supplementary Table 2), which is in agreement with previous reports (Le Tallec et al., 2013). Interestingly, this is true both for control and APH treated cells (Figure 4a). It should be noted that the averaged RT of these genes was mid-S phase, as observed for the fragile genes in 2q22 and 22q12 (Figures 2 and 3). Furthermore, the analysis showed that the only genes whose RT profile was affected by APH treatment were >230kb (Figure 4a). Therefore, we next investigated the association between gene size and APH-induced RT delay. We sub-grouped the large genes (>230 kb) according to size into 4 groups. RT analysis showed that the larger the genes are the later they are replicated both in control and APH treated cells (Figure 4b). Moreover, this analysis confirmed that the RT of large genes is delayed by APH treatment (Figure 4b), implying that the APH-induced RT delay is dependent on gene size but restricted to large genes.

**Figure 4.**
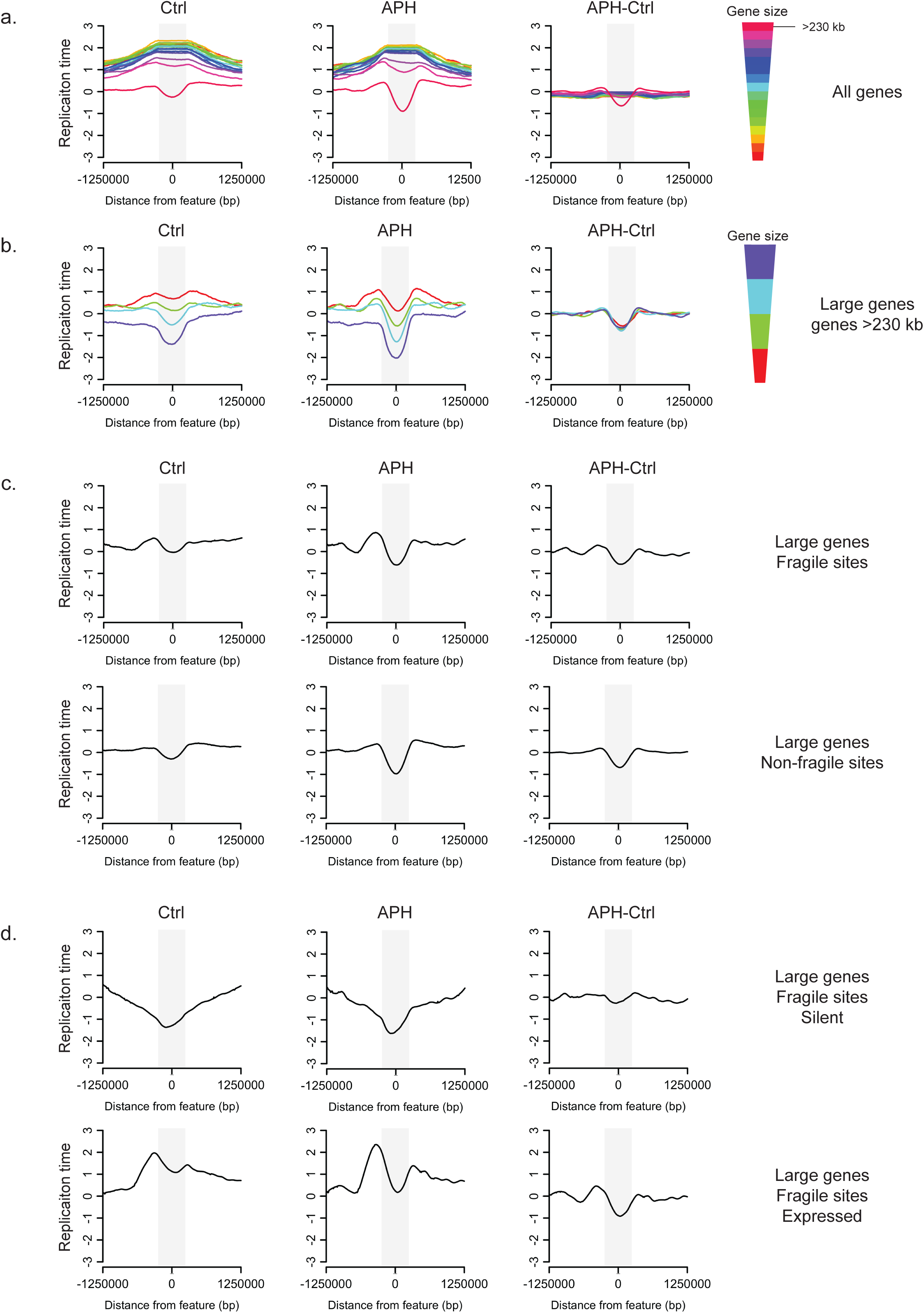
APH delays the RT of large genes. (**a**) Averaged RT of all genes clustered by size in Ctrl and APH treated cells. The difference in RT following APH treatment is presented as the subtraction of RT in Ctrl minus RT in APH treated cells (APH-Ctrl). The Gene body is marked in a grey box and 1 Mb upstream and downstream the genes are presented. Genes are sorted according to size from top to bottom (largest to smallest, respectively) and color coded by cluster as indicated. **(b)** Averaged RT of large genes (> 230 kb) are represented as in (**a**). Genes are sorted according to size from top to bottom (largest to smallest, respectively) and color coded by cluster as indicated. **(c)** Averaged RT of large genes within CFSs (at the top) and in non-fragile regions (at the bottom); in Ctrl, APH and APH-Ctrl. (**d**) Averaged RT of large genes within CFSs. At the top are silent genes, at the bottom are expressed genes, in Ctrl, APH and APH-Ctrl.

Next, we investigated the relationship between delayed replication of large genes and chromosomal instability. We divided the large genes into two groups according to whether or not they overlap a CFS in BJ-hTERT cells. Analysis showed that the average RT of large genes is mid-S phase for both groups of genes (Figure 4c). Moreover, the RT of these large genes is further delayed by APH treatment compared to the RT in control cells (Figure 4c). The delay in RT was also found for both groups of genes, implying that delayed replication of large genes may be necessary but is insufficient to drive chromosomal instability under stress. Next, we analyzed the RT of all the genes in the genome according to whether or not they overlap a CFS in BJ-hTERT cells in order to investigate whether the RT of smaller genes may also be affected by APH treatment within CFSs. RT analysis showed that small genes (<230 kb) are early replicating both in control and APH treated cells, and they are not affected by APH treatment (Supplementary Figure 1). Thus, RT-delay is restricted to large genes whether or not they overlap a CFS in BJ-hTERT cells.

### APH-induced replication delay of large genes is associated with transcription

Previously, expression of large genes has been suggested to associate and promote chromosomal instability at several CFSs (Helmrich et al., 2011; Wilson et al., 2015). However, a recent study reported that the expression level of very large genes affects their stability under replication, suggesting too little or too much transcription is not associated with fragility (Blin et al., 2019). Therefore, we next investigated the expression level of genes following APH treatment, focusing on large genes with delayed replication. For this we used Bru-seq, which maps nascent RNA transcripts, and allows one to measure active transcription rather than steady state transcript levels (Paulsen et al., 2014). The analysis showed that APH treatment did not affect the general transcription program, as only 9 genes were differentially expressed between normal and APH treated cells (Supplementary Table 3). Next, we analyzed the RT profile of large genes overlapping a CFS in BJ-hTERT cells, divided into two groups according to the transcription profile (expressed or silent). The analysis showed that the average RT of silent large genes is late S-phase in control and APH treated cells and their RT is not affected by APH treatment (Figure 4d). Interestingly, the average RT of expressed large genes is early S-phase and their RT is delayed by APH treatment compared to control cells (Figure 4d). In light of the marked difference in the RT profiles of expressed and silent large genes, we next examined the RT of all genes by their transcription profile. The analysis showed that expressed genes are earlier replicating compared to silent genes, as previously reported (Hiratani et al., 2008; Lubelsky et al., 2014; Rivera-Mulia and Gilbert, 2016) (Supplementary Figure 2a). Moreover, large genes showed the largest difference in RT between silenced and expressed genes, from late-S to early-S, respectively, while the RT of the rest of the genes differed within early S-phase (Supplementary Figure 2a). Interestingly, APH-induced delayed RT was found only in expressed large genes (Supplementary Figure 2a). Overall, the RT of large genes seems to be more adjustable and correlated with transcription than the RT of smaller genes.

Next we analyzed the RT of large genes divided into quartiles by the expression level. The results showed that large genes moderately (q3) or highly (q4) expressed were delayed by APH (Supplementary Figure 2b), implying that the RT of large genes with expression levels above the median (q3 and q4) is delayed. In contrast, the RT profile of low expressed large genes (q2) resembled silent genes (q1) (Supplementary Figure 2b). Altogether, these results suggest that for large genes that are transcribed above the median level, the time of replication and their susceptibility to replication stress are transcription dependent.

### Delayed RT of expressed large genes is associated with chromosomal fragility

In light of the transcription-associated RT delay we observed for large genes (Figure 4), we set to explore the expression profile of the large genes within the mapped CFSs 2q22 and 22q12 (Figure 2,3). There are five large genes in 2q22: *ARHGAP15, GTDC1, LRP1B, TEX41* and *THSD7B*, but only the region of *GTDC1* was found to be fragile by FISH mapping (Figure 2). The RT of the *GTDC1* locus is delayed by APH treatment (Figure 2a) and expression analysis showed it is moderately expressed, both in control and APH treated cells (Figure 2b). Interestingly, three of the large genes (*ARHGAP15, TEX41* and *THSD7B*) are silent but their RT is affected by APH treatment and advanced compared to control cells (Figure 2a). In addition, *LRP1B* (1.9 Mb) is a weakly expressed large gene, which is not fragile as indicated by FISH mapping (Figure 2c,d). *LRP1B* is replicated in mid-late S phase in control cells, and it is not delayed by APH treatment. On the contrary, the RT of this gene is advanced in APH treated cells compared to control cells (Figure 2a), indicating that weakly expressed large genes are not delayed by APH treatment. Thus, these results suggest that large genes expressed above the median gene expression level are sensitive to replication stress which leads to fragility.

We further tested this conclusion in the fragile site at 22q12, in which the core fragility was mapped (Figure 3). There are three large genes at 22q12: *SYN3* and *LARGE1*, which according to the FISH mapping are in the non-fragile region and *TTC28* within the core fragility region (Figure 3). The RT of *TTC28* is delayed by APH treatment from early-mid S-phase in control cells to mid-late S in APH treated cells (Figure 3). Expression analysis showed *TTC28* is highly expressed in control and in APH-treated cells with no marked difference in the expression level (Figure 3b). In contrast to *TTC28, SYN3* is replicating in early S-phase in control cells, but it is transcriptionally silent and its RT is not affected by APH treatment (Figure 3). Accordingly, *LARGE1* (∼650 kb), which is not located at the fragile region, is a low expressed large gene (q2). It is replicated in late S-phase both in control and APH treated cells (Figure 3a), and it is not affected by APH treatment. Altogether the results of this analysis show that not all expressed large genes are delayed by APH. Overall, these results support a model by which chromosomal fragility under replication stress is induced at large genes, expressed above the median level, with delayed RT. In control cells the forth mentioned highly expressed large genes are replicating in early-mid S-phase and upon replication stress they are dramatically delayed to mid-late S-phase. The induced V-shaped replication may indicate that under replication stress these regions are origin deficient, thus rendering the few replication forks replicating the genes susceptible to potential obstacles such as active transcription. Our model suggests a novel fragility signature shedding light on the importance of combined contribution of several factors in promoting chromosomal instability.

### Novel fragile sites identified by the fragility signature

The limited resolution of cytogenetic mapping using G-banded metaphase chromosomes may lead to underestimation of chromosomal fragility. Therefore, we set to utilize and test our suggested fragility signature to identify potential novel fragile sites. The fragility signature which is composed of delayed RT specifically of large and highly transcribed genes was identified in 20/24 CFSs we have previously cytogenetically mapped in these BJ cells (Miron et al., 2015), including 2q22 and 22q12 where FISH mapping identified the core fragility region (Figure 2,3). However, a genome-wide screen seeking potential fragile sites based on the existence of large transcribed gene with delayed RT identified additional 93 genes (Supplementary Table 4). Most of these genes are not located within the cytogenetically mapped fragile sites in these cells (Miron et al., 2015). Thus, we set to test whether these genes could indeed be novel fragile genes. To do so, we FISH mapped two candidate fragile genes located outside of fragile bands: IMMP2L (∼890 kb) located at 7q31 and NLGN1 (∼890 kb) located at 3q26 (Figure 5). Both genes replicate in mid-S in control cells and are delayed by APH treatment generating a novel steep V-shaped RT profile (Figure 5a). These genes are also expressed in control and APH treated cells (Figure 5b). Metaphase spreads analysis using FISH probes identified chromosomal breaks at both sites (Figure 5c,d), suggesting that these genes are indeed fragile under replication stress that were missed in the cytogenetic mapping. The results point to the limitation of cytogenetically mapping CFSs and call for revised examination of chromosomal instability across different cell types and in the context of cancer development. Importantly, these results support our proposed fragility signature model emphasizing the importance of a combination of factors impeding DNA replication.

**Figure 5.**
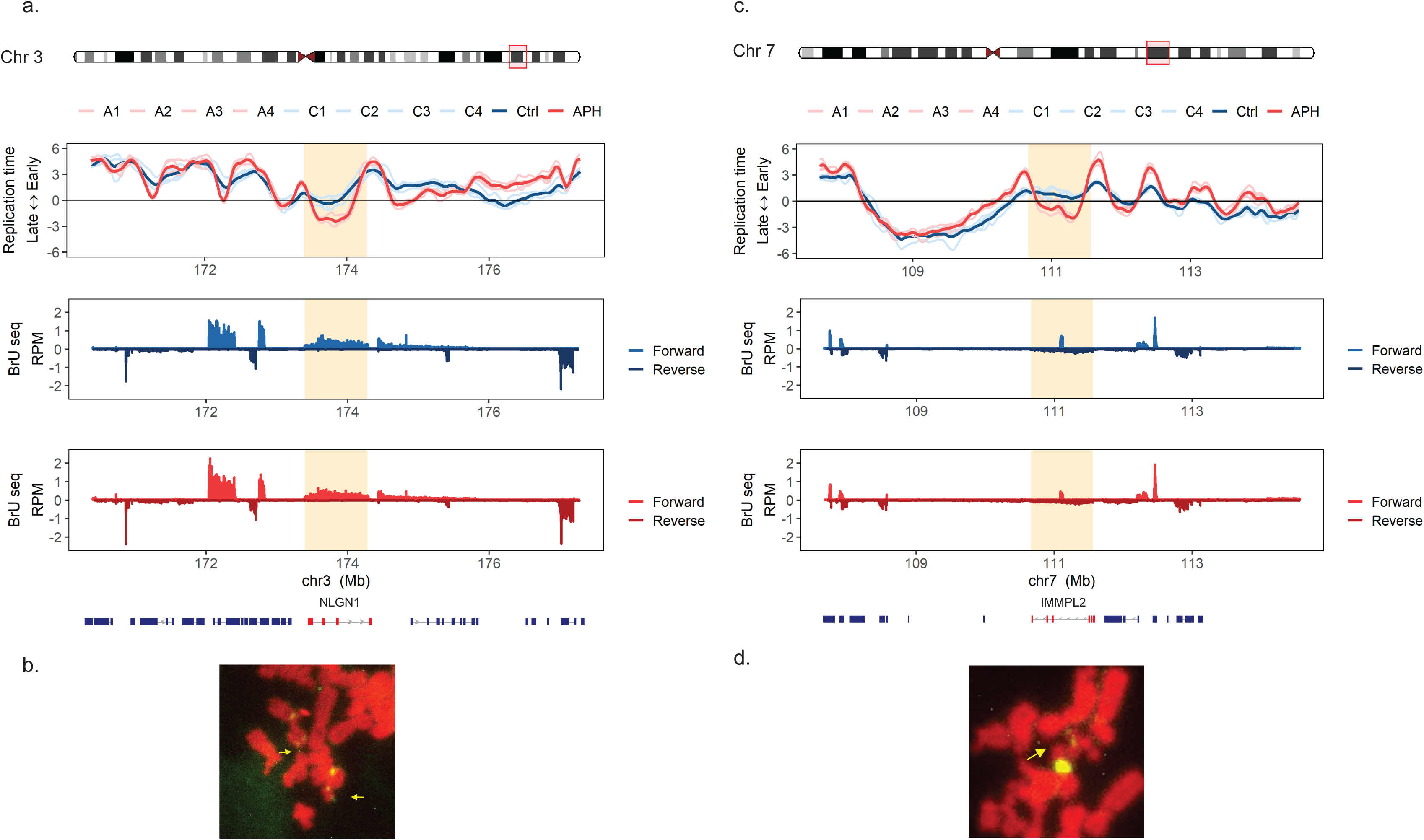
Identification of novel fragile sites. (**a**) RT and nascent RNA transcription of chromosome 3q26. RT and transcription data representation as in Figure 2. Four replicates per condition are presented in RT data color coded according to the legend (Ctrl in blue, APH in red). One replicate per condition is presented in nRNA-seq data, color coded: Ctrl in blue, APH in red. Annotated genes are shown at the bottom. Yellow box marks *NLGN1* gene, which is delayed by APH and expressed both in Ctrl and APH treated cells. (**b**) A representative image of FISH metaphase spread analysis. Yellow arrows indicate chromosomal breaks. (**c**) RT and nascent RNA transcription of chromosome 7q31.RT and transcription data representation as in (**a**). Yellow box marks *IMMP2L* gene, which is delayed by APH and expressed both in Ctrl and APH treated cells. **(d)** A representative image of FISH metaphase spread analysis. Yellow arrow indicates chromosomal break.

## Discussion

Here we have identified a signature for chromosomal fragility upon replication stress, comprised of APH-induced delay in replication timing of early/mid-S phase replicating regions within actively transcribed large genes. CFSs were found to be enriched with delayed RT, coinciding with the fragility core. The fragility signature enabled precise mapping of the core fragility region. Furthermore, the signature enabled the identification of novel fragile sites, not detected cytogenetically, highlighting the improved sensitivity of our approach for identifying fragile sites. These results reveal a link between altered DNA replication and transcription of large genes underlying the mechanism of CFS expression.

DNA replication in eukaryotes is a temporally orchestrated and highly organized process (Rivera-Mulia and Gilbert, 2016). The RT program is tissue specific and is associated with transcriptional activity (Rivera-Mulia and Gilbert, 2016). Previous replication dynamics analyses of single DNA fibers showed that APH-induced replication stress leads to replication fork slowing and even stalling, which subsequently activates dormant origins in an attempt to compensate for the perturbed fork progression (Ge et al., 2007). However, a comprehensive analysis of the RT program under replication stress conditions has not been yet studied. Here, we show that replication stress induced by APH treatment alters the temporal order of replication of only a small part of the genome (∼4% of the genome) (Figure 1), which is highly enriched for CFSs. This suggests that dormant origin activation upon APH treatment is sufficient to preserve the normal and scheduled RT program in the bulk of the genome. It is interesting to note that APH advanced the RT of certain genomic regions (Figure 1.b), presumably by earlier origin activation, suggesting that in addition to a local activation of dormant origins in response to replication stress, in some genomic regions there is a regional response to stress that leads to earlier origin activation. It is tempting to speculate that such advanced RT may play a protective role in maintaining chromosomal stability under replication stress as most advanced RT regions are not within the cytogenetically mapped CFSs, and advanced regions within the mapped CFSs are outside the fragility core region (Figure 2,3).

Previous studies investigating chromosomal instability at four of the most frequently fragile CFSs reported that the fragility core within these CFSs corresponded to the latest replicating origin poor regions (Letessier et al., 2011; Le Tallec et al., 2011), suggesting that upon replication stress these regions may fail to complete DNA replication. However, these RT analyses were performed on cells grown under normal growth conditions. The delayed RT we identified in cells grown under replication stress conditions revealed stress induced- V-shaped RT domain probably resulting from slowed fork progression and lack of origin firing. These findings are in agreement with a previous model in which the fragility core at CFSs is origin deficient (Letessier et al., 2011; Le Tallec et al., 2011). However, comparison of the RT under normal growth conditions to that under replication stress revealed that most of the fragile site regions replicate under normal conditions in early/mid S-phase and under replication stress their RT is delayed. This suggests that the normal time of replication is not the main factor underlying chromosomal fragility. Indeed, most of late replicating regions under normal conditions are not fragile. Altogether the results indicate, that the sensitivity to replication stress manifested as induced delayed-V-shaped RT profile is the basis of CFS expression.

CFS were found to be enriched with large genes (McAvoy et al., 2007; Le Tallec et al., 2013) and for several CFSs, the expression level of the long gene was shown to associate with fragility (Blin et al., 2019; Helmrich et al., 2011; Wilson et al., 2015). Our analyses revealed that delayed RT in CFSs coincided with large genes, supporting the possibility that upon transcription, hazardous collisions between the replication and transcription machineries may occur disrupting fork progression leading to fragility. Moreover, the RT analysis showed that only the RT of large genes is delayed by APH, as compared to the rest of all annotated genes. Further, separating large genes into transcriptionally silent or active groups showed that silent large genes that are late replicating were not affected by APH treatment. Furthermore, there is a lack of origin activation (V-shaped RT pattern) in these silent genes, yet, their RT profiles were not changed under stress conditions. These regions are chromosomally stable, possibly due to lack of transcriptional obstacles. Analysis of the transcribed large genes expressed above the median expression level (q3 and q4) revealed that they are early replicating and their RT is delayed by APH treatment (Figure 4), suggesting that replication-transcription collisions may have perturbed the replication of these genes leading to fragility (Gaillard and Aguilera, 2016; Sollier and Cimprich, 2015). It must be noted that large genes expressed below the median (q2) were hardly affected by APH, implying a transcriptional threshold for imposing replication alterations. Thus, a combination of high transcription of the large gene and insufficient origin activation in the region is underlying CFS expression upon replication stress.

Using the fragility signature of delayed RT along transcriptional active large genes ∼100 regions were identified as potentially fragile under replication stress, of which 20 were identified as CFS by our cytogenetic fragile site mapping (Miron et al., 2015). Hence, we tested whether these regions were not identified as fragile due to the limited fragility resolution of G-banded metaphase chromosomes. Molecular mapping of two such predicted fragile regions showed that they are indeed fragile under APH treatment (Figure 5). These results highlight the improved sensitivity of our approach for identifying fragile sites, overcoming the limitations of cytogenetic mapping. Furthermore, as fragile sites are cell type specific (Hosseini et al., 2013; Murano et al., 1989; Le Tallec et al., 2013) revised mapping of fragile sites using RT profiling in cells grown under normal compared to replication stress conditions, should be performed for identification of the entire repertoire of fragility in different cell types.

Altogether, our results show that fragility is affected by the combined effect of replication and transcription driving chromosomal instability under replication stress. Cancer development is promoted by replication–induced genomic instability (Halazonetis et al., 2008; Macheret and Halazonetis, 2015), and changes in gene expression (Bradner et al., 2017). Thus, investigating the RT changes in cancer cells may contribute to our understanding of genomic instability in cancer.

## Methods

### Cell culture and treatments

Human foreskin fibroblasts BJ-hTERT cells (Clontech) were grown in DMEM supplemented with 10% fetal bovine serum, 1000,000 U l^−1^ penicillin and 100 mg l^−1^ streptomycin. The cells were tested and found negative for mycoplasma. Cells were treated with 0.2 μM Aphidicolin and 1.46 mM caffeine in growth media for 24 hours prior to fixation.

### FISH mapping on Metaphase chromosomes

Cells were treated with 100 ng ml−1 colcemid (Invitrogen, Carlsbad, CA, USA) for 40 min in incubator at 5% CO2. Then, cells were collected by trypsinization, treated with hypotonic solution at 37 °C for 30 min and fixed with multiple changes of methanol:acetic acid 3:1. Fixed cells were kept at −20 °C until analysis. For analysis of total gaps, constriction or breaks chromosomes were stained with propidium iodide and blindly analyzed. For fluorescent in situ hybridization (FISH) analysis, BAC clones (RP11-316D5, RP11-265L20, RP11-158G12, RP11-61D21, RP11-707K3, RP11-354I1, RP11-706F7) were labeled with by nick translation. In order to evaluate metaphases which experienced similar level of replication stress, metaphases that exceeded a threshold of 10 breaks were not included in the FISH evaluation.

### Genome-Wide RT Profiling

Genome-wide RT profiles were constructed as previously described (Marchal et al., 2018). Briefly, cells were pulse labeled with BrdU, separated into early and late S-phase fractions by flow cytometry, and processed by Repliseq. Sequencing libraries of BrdU-substituted DNA from early and late fractions were prepared by NEBNext Ultra DNA Library Prep Kit for Illumina (E7370; New England Biolabs). Sequencing was performed on an Illumina-HiSeq 2500 sequencing system by 50-bp single-end reads. Reads with quality scores above 30 were mapped to the Hg38 reference genome using bowtie2. Approximately 8 million uniquely mapped reads were obtained from each library. Read counts were binned into 5-kb nonoverlapping windows, and log2 ratios of read-counts between early and late fractions were calculated. Plots of RT profiles were generated in R project for Statistical Computing (http://www.r-project.com). Averaged RT profiles of genes clustered by size were generated using genmat package for R (https://rdrr.io/github/dvera/genmat/).

### Clustering Analysis

RT profiles were expressed as numeric vectors. Nonoverlapping and variable regions (100-kb windows) were defined as those with differences ≥1 in pairwise comparisons between all samples. Unsupervised hierarchical and kmeans clustering analysis were performed using Cluster 3.0 (de Hoon et al., 2004). Heatmaps and dendrograms were generated in R project for Statistical Computing (http://www.r-project.com).

### Bru-seq

Bru-seq was performed as previously described. Briefly, bromouridine (Bru) (Aldrich) was added to the media to a final concentration of 2 mM and incubated at 37°C for 30 min. Total RNA was isolated using TRIzol reagent (Invitrogen), and Bru-labeled RNA was isolated by incubation of the isolated total RNA with anti-BrdU antibodies (BD Biosciences) conjugated to magnetic Dynabeads (Invitrogen) under gentle agitation at room temperature for 1 h. cDNA libraries were prepared from the isolated Bru-labeled RNA using the Illumina TruSeq library kit and sequenced using Illumina HiSeq sequencers at the University of Michigan DNA Sequencing Core. The sequencing and read mapping were carried out as previously described (Paulsen et al., 2014).

Reads were mapped to hg38 using bowtie. RPM-normalized read densities were calculated with bedtools2 (Quinlan and Hall, 2010) genomecov, using a scaling factor of 1000000/(number of parsed reads in library). The profiles are shown at 1kb window bins.

## Supplementary Figures

**Supplementary Figure 1. Large genes are delayed by APH irrespective to fragile site location**. Averaged RT of all genes within CFSs (a) and in non-fragile regions (**b**) clustered by size in Ctrl and APH treated cells. The difference in RT following APH treatment is presented as the subtraction of RT in Ctrl minus RT in APH treated cells (APH-Ctrl). The Gene body is marked in a grey box and 1 Mb upstream and downstream the genes are presented. Genes are sorted according to size from top to bottom (largest to smallest, respectively) and color coded by cluster as indicated.

**Supplementary Figure 2. Delayed RT of large genes is transcription dependent**. (**a**) Averaged RT of all genes clustered by size and expression profile: silent at the top and expressed at the bottom, in Ctrl and APH treated cells. The difference in RT following APH treatment is presented as the subtraction of RT in Ctrl minus RT in APH treated cells (APH-Ctrl). The Gene body is marked in a grey box and 1 Mb upstream and downstream the genes are presented. Genes are sorted according to size from top to bottom (largest to smallest, respectively) and color coded by cluster as indicated. (**b**) Averaged RT of large genes (> 230 kb) binned by expression quartile 1-4, where q1 is lowest expression and q4 is the highest expression level.

## Supplementary Tables

**Supplementary Table 1. RT signatures**. List of genomic regions within each RT signature. 1277 windows of 100kb identified as RT variable between Ctrl and APH treated cells are presented.

**Supplementary Table 2. Averaged RT profiles of genes**. List of averaged RT profiles for genes clustered into 16 groups according to size. Genes shorter than 5kb were filtered out of the analysis.

**Supplementary Table 3. APH induced differentially expressed genes**. List of genes differentially expressed, either down- or up-regulated by 0.2μM APH treatment for 24 hours, as detected by Bru-seq.

**Supplementary Table 4. Fragility signature candidate genes**. List of candidate fragile genes according to the fragility signature: expressed large genes, delayed under APH treatment.

