## Supplementary figures and images for "Replication Timing and Transcription Identifies a Novel Fragility Signature Under Replication Stress"

### Supplementary Figures

#### Supplementary Figure 1

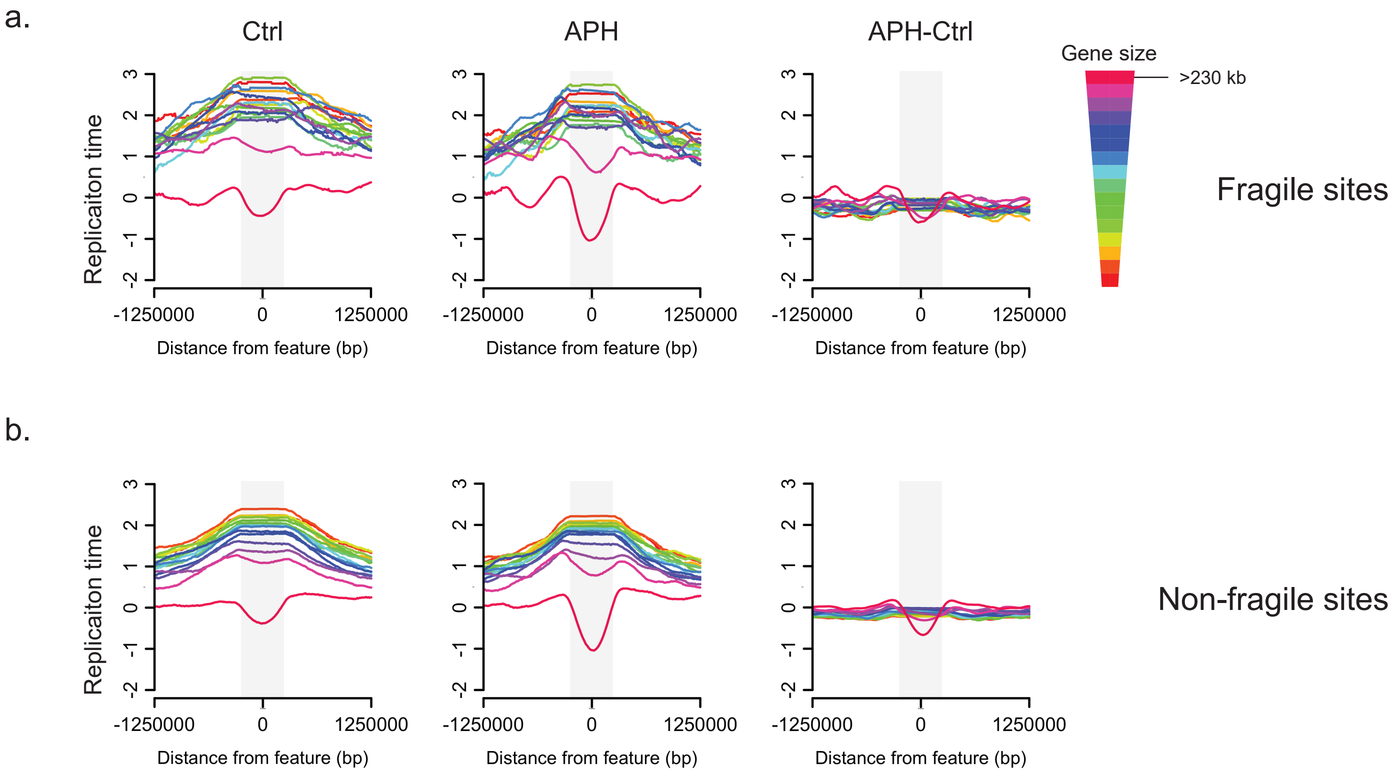

#### Supplementary Figure 2

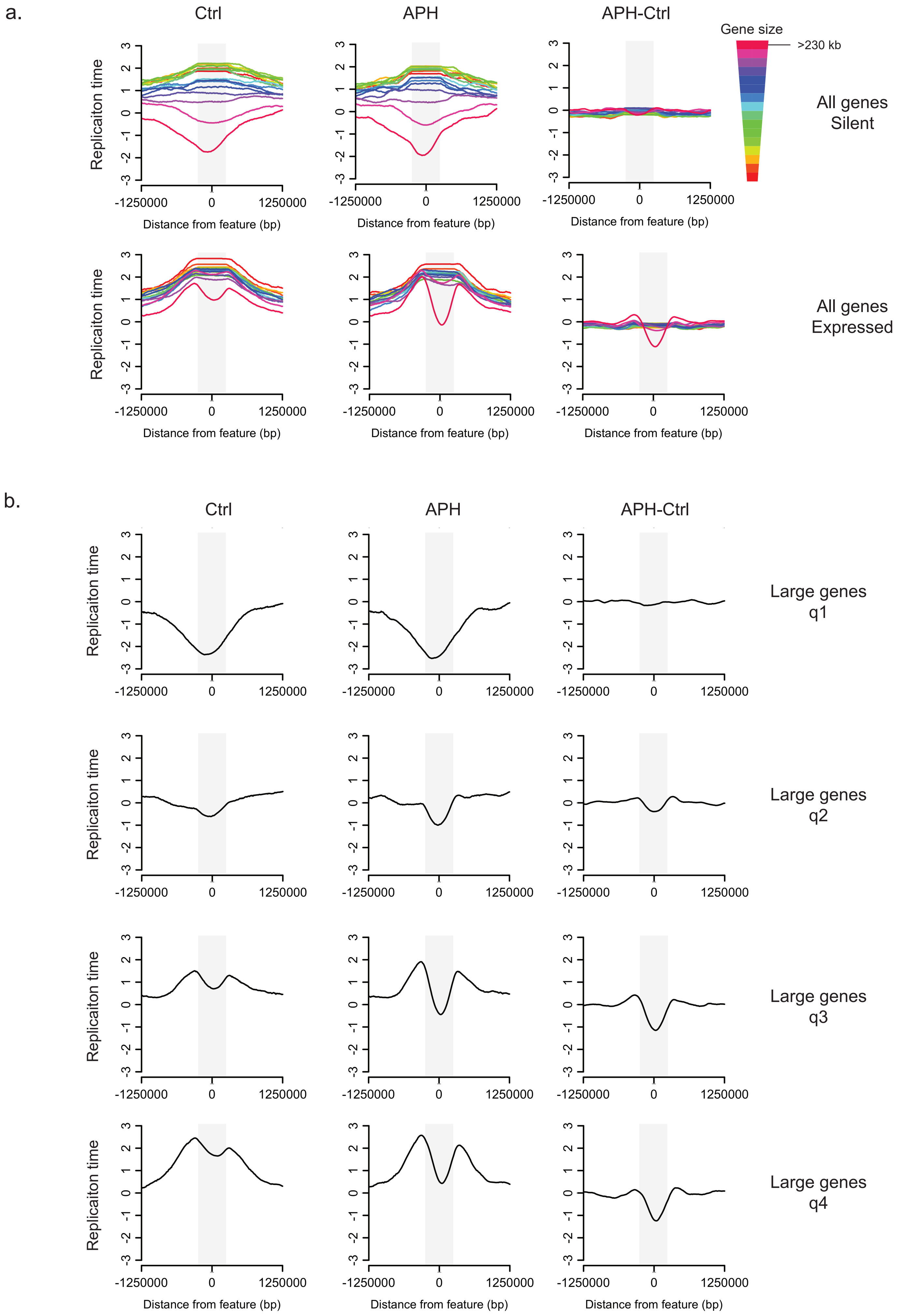
